# Dysregulation of the engulfment pathway in the gut fuels Inflammatory Bowel Disease

**DOI:** 10.1101/280172

**Authors:** Katherine Suarez, Eileen Lim, Sujay Singh, Matheus Pereira, Linda Petronella Joosen, Stella-Rita Ibeawuchi, Ying Dunkel, Yash Mittal, Samuel B. Ho, Ranajoy Chattopadhyay, Monica Guma, Brigid S. Boland, Parambir S. Dulai, William J. Sandborn, Pradipta Ghosh, Soumita Das

## Abstract

**BACKGROUND & AIMS:** Luminal dysbiosis is ubiquitous in inflammatory bowel disease (IBD), but how the microbes trigger pro-inflammatory cascades in the epithelial and phagocytic cells remains unknown. Here we investigated the role of the microbial sensor ELMO1 (Engulfment and Cell Motility Protein-1) in sensing and responding to IBD-associated microbes in the gut epithelium and in macrophages.

**METHODS:** A stem cell-based technique is used to grow enteroids from WT and ELMO1^−/−^mice and from colonic biopsies of patients with IBD and subsequently differentiate them into enteroid-derived monolayers (EDMs) that mimic the gut epithelium/Gut in a dish. EDMs infected with IBD-associated invasive *E. coli*-LF82 were analyzed for bacterial internalization, cytokine production and monocyte-recruitment when co-cultured with monocytes.

**RESULTS:** Expression of ELMO1 is elevated in the colonic epithelium and in the inflammatory infiltrates within the lamina propria in IBD, higher expression correlated with elevated expression of pro-inflammatory cytokines, MCP-1 and TNF-α. ELMO1-/-murine EDMs displayed a significant reduction of bacterial internalization through epithelial tight junctions and in MCP-1 production compared to WT mice. MCP-1 that is released from the epithelium recruited monocytes. Once recruited, macrophages required ELMO1 to engulf the bacteria and propagate a robust pro-inflammatory cytokine storm (TNF-α).

**CONCLUSIONS:** ELMO1 couples microbial-sensing to inflammation in both phagocytic and non-phagocytic host cells; it is required for the production of MCP-1 in the epithelium and TNF-α in macrophages. Findings raise the possibility that upregulation of epithelial ELMO1 and the epithelial ELMO1→MCP-1 axis may serve as an early biomarker and therapeutic target, respectively, in IBD and other disorders of inflammation.

## Introduction

Inflammatory bowel disease (IBD) affects 1.5 million people in the United States. IBD is a multifactorial disease where both environmental and genetic factors are involved ^1, 2^. Current treatment consists of corticosteroids, immunosuppressants, biologics, and in some cases, surgery, but no treatments are universally effective^3^. Thus, there is an urgent need to further understand the pathogenesis of IBD, in order to develop new therapeutics. One potential therapeutic approach is the detection and targeting of specific bacteria-mediated inflammation that initiates in the earlier stages of the disease and present in chronic infection. Microbial dysbiosis is one of the critical component of IBD^4-7^. IBD samples contained abnormal GI microbiotas, a shift in the microbiota toward more pro-inflammatory species characterized by depletion of commensal bacteria, notably members of the phyla Firmicutes, Bacteroidetes, *Bifidobacterium, Clostridia* and enrichment of *Actinobacteria* and *Gamma proteobacteria*^8-10^. Adherent-invasive *E. coli* (AIEC) is the major pathovar of the species *Escherichia coli* that has isolated from CD patients^11^.

Intestinal dysbiosis triggers distinct steps at the host-microbe interface--First, recognition and internalization of pathogen by phagocytic receptor/mediator controls the downstream inflammatory responses^12, 13^. For example, *AIECs* must adhere and invade the gut epithelial cells, followed by engulfment to macrophages and induction of inflammatory pathways in host cells^14^. A finite inflammatory response is essential to clear the infection without extensive collateral damage and destruction of host tissue. However, the defective clearance of bacteria is the hallmark in CD, with a resultant retention of the engulfed bacteria within macrophages propagates inflammation, creates a vicious cycle of subsequent rounds of recruitment of monocytes and T lymphocytes and the release of proinflammatory cytokines. The underlying mechanism of host epithelium, which is breached by invading pathogens and initiates responses to mount an inflammatory cytokine storm is unclear.

We recently demonstrated that Engulfment and cell motility protein 1 (ELMO1) is a microbial sensor that enables macrophages to engulf enteric bacteria, and coordinately mount inflammation while orchestrating bacterial clearance via the phagolysosomal pathway^15-17^. ELMO1 binds the Pattern Recognition Receptor (PRR) Brain Angiogenesis Inhibitor-1 (BAI1) which recognizes bacterial Lipopolysaccharide (LPS)^18^. BAI1-ELMO1 signaling axis activates Rac1 and induces pro-inflammatory cytokines Tumor necrosis factor-α (TNF-α) and monocyte Chemoattractant protein 1 (MCP-1)^15, 19^. BAI1-ELMO1 signaling axis regulates the expression of ATP-binding cassette transporter ABCA1 which is linked to the development of cardiovascular diseases (CVD)^20^. Genome wide association studies (GWAS) have revealed the association of single nucleotide polymorphisms (SNPs) in ELMO1 with IBD, rheumatoid arthritis (RA) kidney disease and diabetic nephropathy^21-23^. ELMO1 is required for the induction of several proinflammatory cytokines that are known to drive a plethora of inflammatory diseases including IBD, CVD and RA^15^. Among them, MCP-1 is a chemokine, which plays a significant role in the recruitment of mononuclear cells to the site of inflammation; it is also one of the major cytokines involved in inflammatory diseases like IBD^24^.

The function of ELMO1 in phagocytic cells is clear; however, its role in the non-phagocytic cells, i.e., the gut epithelium remains unknown. We hypothesized that the sensing of IBD-associated microbes by ELMO1 in the gut epithelium, the first line host defense, may serve as an upstream trigger for immune cell-mediated cytokine storm. We used the cutting-edge stem cell based-enteroids as the model system to interrogate the role of ELMO1 in epithelial cells faced with the dysbiosis in CD patients, and defined a specific need for the engulfment pathway in the induction of MCP-1. The generation of MCP-1 by the epithelium appears to be followed by monocyte recruitment at the site of inflammation. Subsequently, bacteria enter monocytes in an ELMO1-dependent manner and trigger the release of TNF-a, thereby, propagating the choronic inflammatory cascade that is the hallmark of IBD. Findings raise the possibility that targeting ELMO1 may help simultaneously blunt both the ELMO1-MCP-1 and the ELMO1-TNF-α signaling axes in the epithelium and the macrophages, respectively, to combat the inflammation in IBD.

## Materials and Methods

All authors had access to the study data and reviewed and approved the final manuscript.

### Bacteria and Bacterial Culture

Adherent Invasive *Escherichia coli* strain LF82 (*AIEC-*LF82), isolated from the specimens of Crohn’s disease patient, was obtained from Arlette Darfeuille-Michaud^11^. A single colony was inoculated into LB broth and grown for 8 h under aerobic conditions and then under oxygen-limiting conditions. Cells were infected with a multiplicity of infection (moi) of 10.

### Animal

WT and ELMO1 KO C57BL/6 mice were gender- and age-matched litter mate, that used to isolate intestinal crypts and macrophages. Animals were bred, housed, and euthanized according to all University of California San Diego Institutional Animal Care and Use Committee (IACUC) policies.

### Human Subjects

All the IBD patients were seen at the University of California, San Diego at the IBD-Center. They were recruited and consented using a study proposal approved by the Institutional Review Board of University of California, San Diego. The clinical phenotype and information were assessed by an IBD specialist.

### Cell Lines and Cell Culture

Control and ELMO1 small-hairpin RNA (shRNA) macrophage (J774) cells were maintained in high glucose Dulbecco’s modified Eagle’s medium (DMEM) containing 10% fetal calf serum, as described previously^15^. THP-1 cells were maintained in RPMI media containing 10 mM HEPES, 10 % FBS at a cell concentration of lower than 10^6^cells/ml^19^.

### Isolation of enteroids from colonic specimens of mouse and human

Intestinal crypts were isolated from the colonic tissue specimen by digesting with Collagenase type I [2 mg/ml; Invitrogen] and cultured in stem-cell enriched conditioned media with WNT 3a, R-spondin and Noggin ^25-27^. The details are mentioned in the Supplementary Materials and Methods.

### The preparation of Enteroid-derived monolayers (EDMs)

To prepare EDMs, single cells from enteroids in 5% conditioned media was added to diluted Matrigel (1:30) as done before^28^. In some cases, the EDMs were also differentiated for 2 days in advanced DMEM/F12 media without Wnt3a but with R-spondin, Noggin, B27 and N2 supplements and 10 mM ROCK inhibitor^26^. As expected, this results in a marked reduction in the expression of the stemness marker Lgr5 in EDMs^26^.

### Isolation of murine intestinal macrophages

Intestinal macrophages were isolated as described previously^15, 17, 18^ and in the supplementary Materials and Methods

### Bacterial internalization by gentamicin protection assay

Approximately 2×10^5^cells were plated onto a 0.4 μm pore transwell insert and infected with bacteria with moi 10. Bacterial internalization was determined after 6 h of infection of WT and ELMO1-/-EDMs with *AIEC-*LF82 by gentamicin protection assay^15, 18^.

### Monocyte Recruitment Assay

WT and ELMO1-/-EDMs were plated in the transwell for polarization, and infected with *AIEC-*LF82 for 6 h. Supernatant from the basolateral chamber was collected and placed in the new 24-well plate, and 6.5 mm 8-μm pore-sized transwells (Costar) where THP-1 cells or peripheral blood-derived monocytes in OptiMEM (Gibco) were placed on the apical chamber. In another assay, the EDM layer was collected and flipped and placed on the bottom of the transwell. The number of recruited live monocytes were measured after 1, 2, 8, 16 and 24 h.

### Cytokine Assays

Supernatants were collected from the basolateral chamber either uninfected or after infected cells. MCP-1 was measured using the Mouse CCL2 (MCP-1) ELISA Ready-Set-Go Kit according to manufacturer’s instructions (eBioscience). Supernatants were collected from control or ELMO1-depleted J774 cells after *AIEC-*LF82 infection and TNF-a was measured using the ELISA kit from BD bioscience.

### RNA Preparation, Real-Time Reverse-Transcription Polymerase Chain Reaction

EDM layer following infection with *AIEC-*LF82 was collected for RNA isolation followed by quantitative RT-PCR as described in details in the Supplementary Materials and Methods.

### Immunohistochemistry

A total of 8 colonic specimens of known histologic type (3 normal colorectal tissue; 3 ulcerative colitis and 2 Crohn’s disease) were analyzed by IHC using anti-ELMO1 antibody (1:20, anti-rabbit antibody from Novus). Briefly, formalin-fixed, paraffin-embedded tissue sections of 4 μm thickness were cut and placed on glass slides coated with poly-L-lysine, followed by deparaffinization and hydration. Heat-induced epitope retrieval was performed using citrate buffer (pH-6) in a pressure cooker. Tissue sections were incubated with 0.3% hydrogen peroxidase for 15 min to block endogenous peroxidase activity, followed by incubation with primary antibodies for 30 min in a humidified chamber at Room temperature. Immunostaining was visualized with a labeled streptavidin-biotin using 3,3′-diaminobenzidine as a chromogen and counterstained with hematoxylin.

### Gene expression analysis

The association between the levels of ELMO1 and MCP-1 (*CCL2*) mRNA expression was tested in a cohort of normal colon tissue as described in details in the Supplementary Materials and Methods.

### Confocal Microscopy

WT and ELMO1-/-EDMs were plated onto 8-well chamber slides (Millicell) and infected with bacteria with moi 10. After infection, cells were stained for LAMP-1 (Lysosomal Associated Membrane Protein-1) and ZO1 (Zonula Occludens) following the protocols as mentioned in the Supplementary Materials and Methods.

### Statistical analysis

Bacterial internalization, monocyte recruitment assays and ELISA results were expressed as the mean ± SD and compared using a two-tailed Student’s t test. Results were analyzed in the Graphpad Prism and considered significant if p values were < 0.05.

## Results

### 3.1. Expression of ELMO1 positively correlates with that of pro-inflammatory cytokines, MCP-1 and TNF-α

Previously we showed that ELMO1 is involved in intestinal inflammation and microbial sensing^15^. To understand the role of ELMO1 in IBD, we analyzed publicly available datasets [Pubmed; Gene Expression Omnibus (GEO) datasets GDS1330 / 24F24]^29^ for expression of ELMO1 in the sigmoid colons of UC and CD patients (Figure 1*A*). In healthy humans, ELMO1 expression is heterogeneous with an average expression of ~0.10 Arbitrary Units (AU). In patients with CD and UC however, expression is higher in both when compared to healthy humans (with *p* value of 0.036).

**Figure 1:**
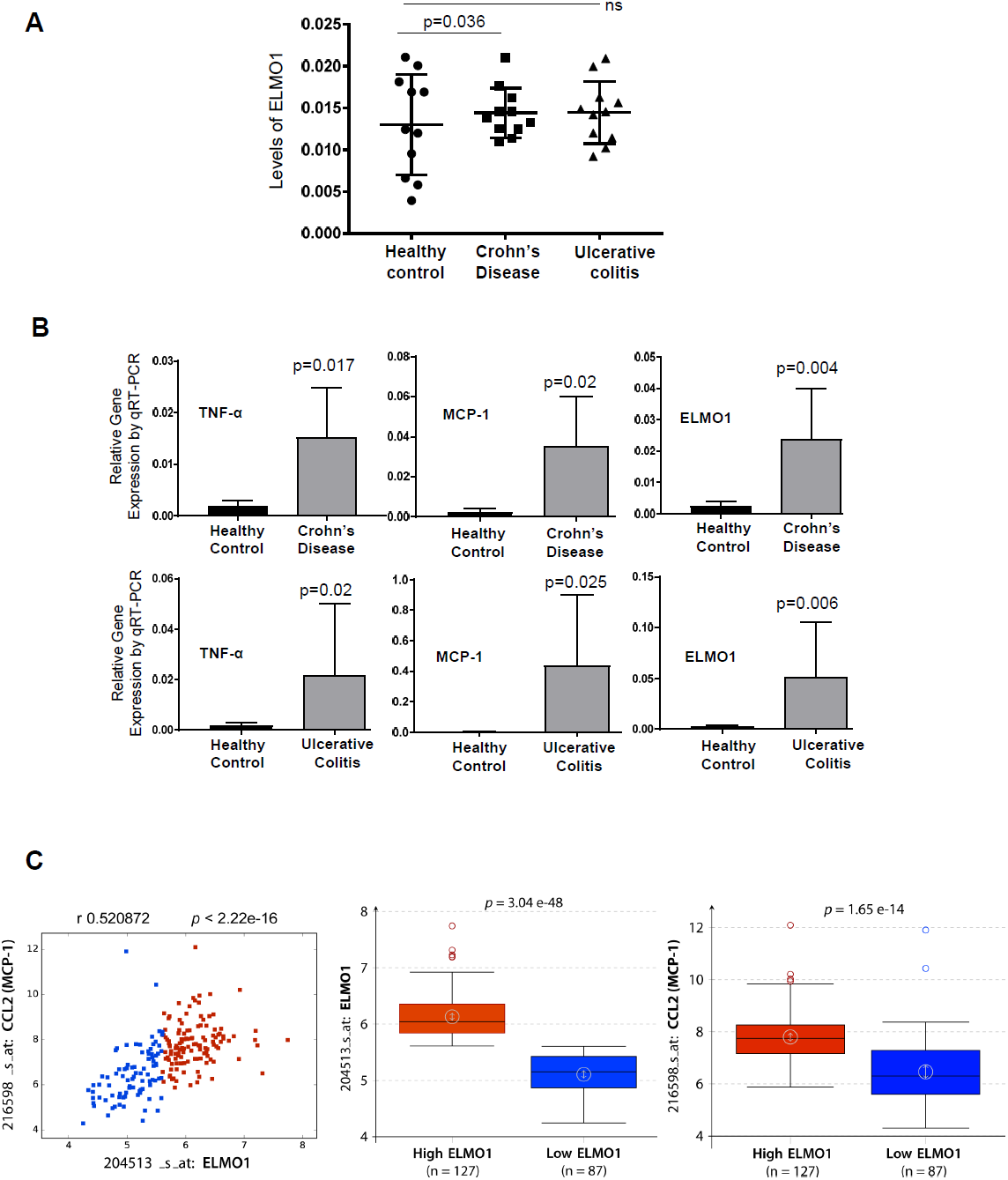
ELMO1 is expressed in the gut epithelium, and its elevated expression in the gut correlates with inflammation. *(A)* Gene Expression Omnibus (GEO) repository was queried for the patterns of expression of ELMO1 in publicly available cDNA microarrays^29^[GDS1330/ GSE 1710; performed using mucosal biopsy samples from sigmoid colons of normal healthy controls (n = 11), patients with Crohn disease (CD; n = 10) and ulcerative colitis (UC; n = 11). Findings are displayed as scatter plots. *(B)* Expression of ELMO1, MCP-1 and TNF-α were determined by qRT-PCR on the RNA isolated from colonic biopsies obtained from healthy controls and patients with Crohn’s disease or Ulcerative colitis (n=6-8 samples/group). *(C)* The association between the levels of ELMO1 and MCP-1 (*CCL2*) mRNA expression was tested in a cohort of 214 normal colon samples available in NCBI-GEO data-series (see *Methods*). *<u>Left</u>*: Graph displaying individual arrays according to the expression levels of *CCL2* and *ELMO1* in 214 normal colon tissues. Probe ID used for each gene is shown. Blue and red indicate samples stratified into high (n = 127) vs low (n = 87) ELMO1 groups using StepMiner algorithm. *<u>Middle</u>*: Box plot comparing the levels of ELMO1 between high vs low ELMO1 groups. *<u>Right</u>*: Box plot comparing the levels of MCP-1 between high vs low ELMO1 groups.

The relative expression of pro-inflammatory cytokines TNF-α, MCP-1, and ELMO1 was assessed in RNA isolated from human biopsy samples from healthy controls or with IBD (Figure 1*B*). Compared to healthy controls, expression of both TNF-α and MCP-1 were elevated ~6-fold in patients with active CD. As for ELMO1, its expression was elevated ~4-fold compared to healthy controls.

The association between the levels of ELMO1 and MCP-1 (CCL2) was studied in the publicly available NCBI-GEO data-series where RNA Seq data from 214 normal colons showed that ELMO1 and MCP-1 genes display a Boolean relationship in which, if the levels of expression of one is high, usually the other is also high (Figure 1*C*). These findings suggest a more fundamental gene expression signature that is conserved despite population variance.

To determine if elevated ELMO1 mRNA levels translate into elevated protein expression, and if so, which cell types contribute to such elevation we performed immunohistochemistry (IHC) on colonic biopsies from healthy controls or patients with UC or CD (Figure 2). We detected ELMO1 both in the epithelium and the lamina propria of the normal gut; expression was variable between individuals. In biopsies from patients with CD or UC, ELMO1 expression was elevated both in the epithelium and the lamina propria, but most strikingly in the diseased epithelium.

**Figure 2:**
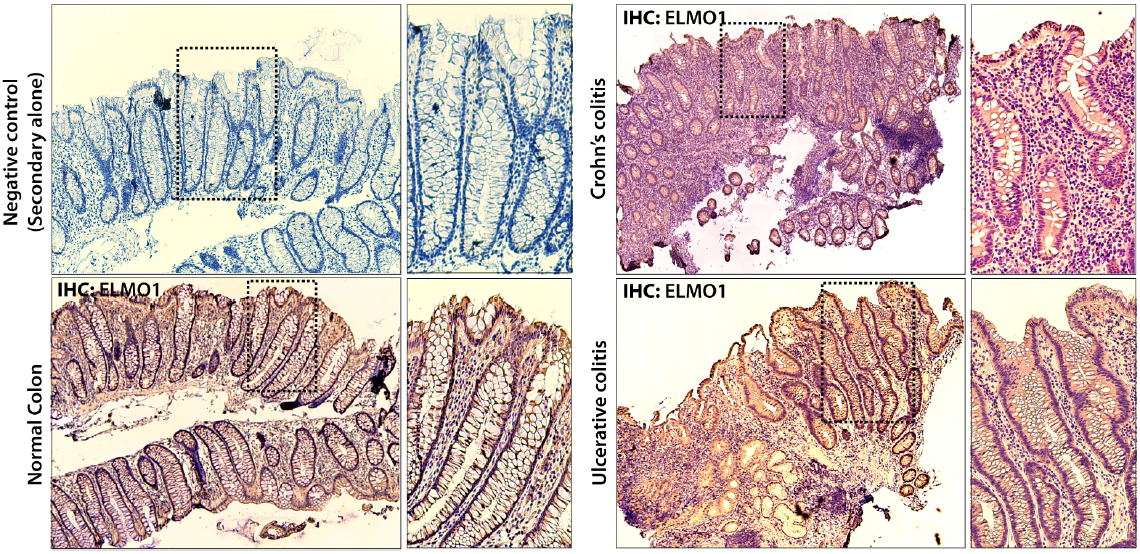
Expression of ELMO1 was determined by IHC on biopsies obtained from healthy controls (normal colon; left) or patients with UC or CD (right). A part of the section was stained with secondary antibody that was shown as negative control.A representative figure was selected from all the sections stained with ELMO1.

Taken together, these findings indicate that expression of ELMO1 is elevated in colons of patients with IBD; expression was detected not just in the lamina propria, presumably contributed by infiltrating immune cells, but also in the epithelial lining.

### 3.2. The adherent-invasive E. coli (AIEC) is a model bacterium to study the role of the host engulfment pathway in CD pathogenesis

To investigate the role of ELMO1 in the IBD-afflicted gut epithelium, we utilized the stem-cell derived 3D intestinal organoids or enteroids^30^. Enteroids mimic the *in vivo* situation with four different cell types’ epithelial, goblet, Paneth cells and enterocytes^31-33^. Therefore, the use of human crypt-derived enteroids recreates normal intestinal physiology. We generated enteroids (Figure 3*A i*) from colonic biopsies obtained from healthy controls and CD patients, and enteroid-derived monolayers (Figure 3*A ii*) (*Table 1* for patient demographics and clinical information).

**Figure 3:**
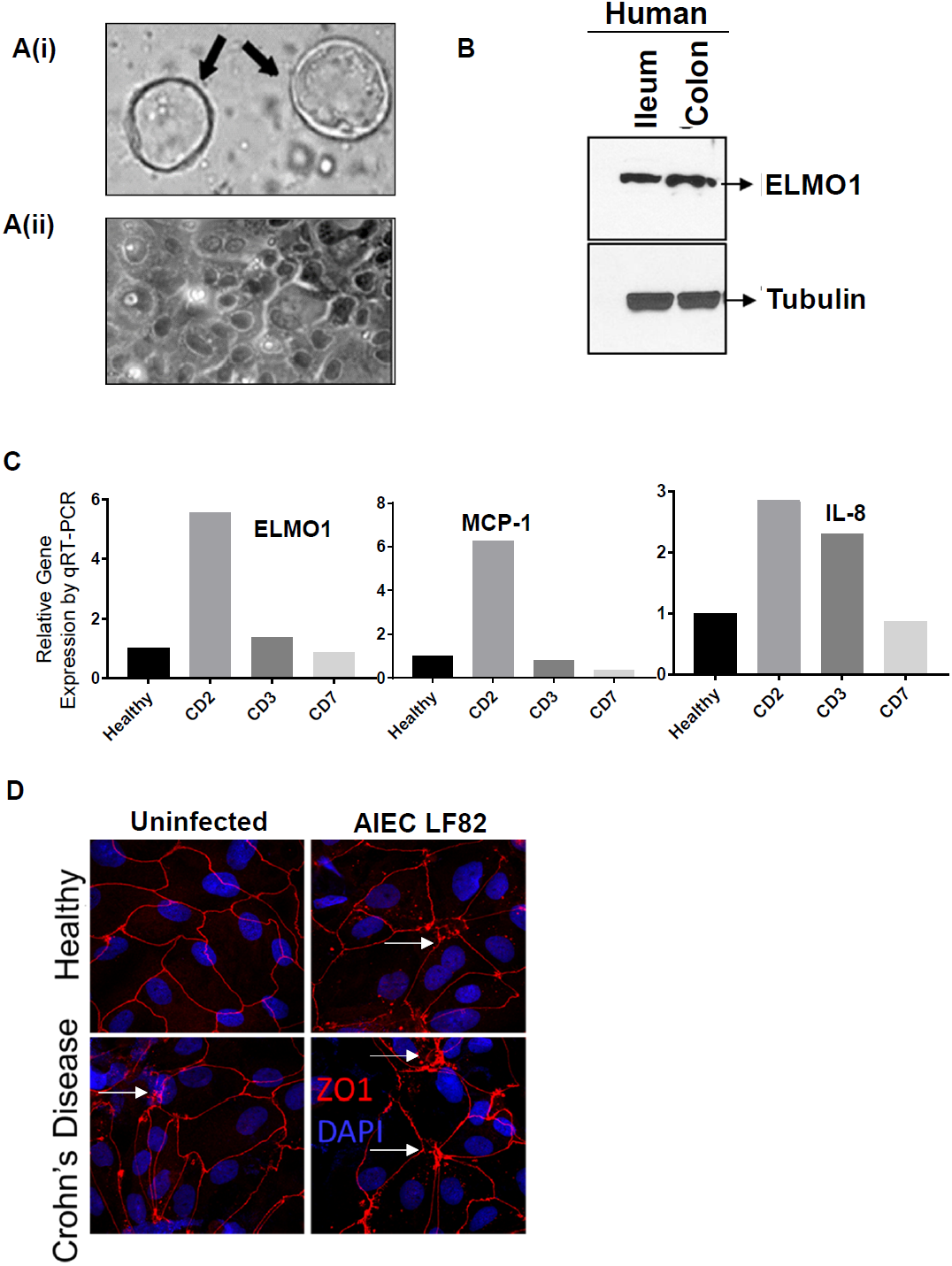
Enteroid-derived monolayers as a model system to selectively interrogate the role of the gut epithelium in CD. *(A)*(i) Enteroids isolated from colonic biopsies that were obtained from either healthy controls or patients with CD were viewed by light microscopy. A representative image of spheroids (arrows) is displayed. (ii) Enteroid-derived monolayers (EDM) prepared from the enteroids via terminal differentiation (see *Methods*) were viewed by light microscopy. A representative image of the EDM is shown. *(B)* The levels of expression of ELMO1 (75 kD) was detected by immunoblotting of enteroids derived from the terminal ileum and sigmoid colon of a representative healthy subject; where a-Tubulin was used as a loading control. *(C)* The expression of ELMO1, MCP-1 and IL-8 were measured in the EDMs isolated from colonic biopsies obtained from one healthy and three CD patients (description in Table 1). Bar graphs display the fold change in expression normalized to the healthy control. *(D)* EDMs derived from colonic biopsies obtained from healthy subjects and from patients afflicted with CD were infected (right) or not (left) with *AIEC*-LF82 prior to fixation and stained for ZO-1 (red), a marker for TJs and nucleus (DAPI; blue). Disruptions in TJs is marked (arrowheads). In healthy EDMs, disrupted TJs are seen exclusively after infection with *AIEC*-LF82 (compare two upper images). In CD-derived EDMs, disrupted TJs were noted at baseline (lower left), almost to a similar extent as after infection with *AIEC*-LF82 (compare two lower images).

Table: Description of the IBD samples

**Table: Description of the IBD samples**
| **Patient#** | **Age/Gender** | **Anatomic location** | **Histopath** | **Treatment history** |
| --- | --- | --- | --- | --- |
| CD2 | 27/Female | Left colon | Colonic mucosa with no diagnostic alteration. No granulomas, viral cytopathic effect or dysplasia identified | Remicade<br>12 days |
| CD3 | 54/Male | Rectum | Colonic mucosa with no diagnostic alteration. | Humira<br>6 days |
| CD7 | 19/Female | Left colon | Severely active colitis with ulceration, granulation tissue, and scattered granulomas | Ustekinumab<br>At the time of biopsy |

First, we confirmed that ELMO1 is indeed expressed in the human enteroids from colon and ileum by immunoblotting (Figure 3*B*) and by quantitative real-time RT-PCR (qRT-PCR; Figure 3*C*). When compared to healthy controls, levels of ELMO1 mRNA were elevated in CD-derived enteroids (Figure 3*C*). Although the degree of elevation, was heterogeneous, as expected given that the patients were receiving anti-TNF-α therapy before biopsies were taken (Table 1), the degree of increase in ELMO1 positively tracked with markers of inflammation, i.e., expression of MCP-1 and IL-8 (Figure 3*C*).

To understand the role of epithelial ELMO1 following exposure to IBD-associated bacteria, we chose to study adherent-invasive *E. coli* (*AIEC*-LF82) as a model bacteria^11,34,35^. Because invasive microbes attack the integrity of epithelial tight junctions (TJs), trigger a redistribution of apical tight junction protein Zonula Occludens-1 (ZO-1)^36^and thereby, breach the epithelial barrier function during invasion, we investigated epithelial TJs between healthy or CD-derived enteroids after exposure to *AIEC*-LF82. TJs were clearly defined and intact in the uninfected healthy EDMs, but they were disrupted when EDMs were infected with *AIEC*-LF82 (Figure 2*D*). In fact, the extent of disruption was almost similar (i.e., ~90-95 % area affected) to uninfected CD-derived EDMs at baseline. Upon infection, the CD-derived EDMs showed increased levels of ZO1 at the TJs, which may be due to a short-term protective mechanism(s) that recruits ZO1 to resist infection/stress-induced TJ collapse. These findings confirm that the CD-associated *AIEC*-LF82 can indeed disrupt epithelia TJs in the healthy epithelium, much like that seen in CD-derived EDMs at baseline, providing a solid rationale for continuing to study the interplay between the *AIEC*-LF82 and epithelial ELMO1.

### 3.3. ELMO1 is required for the engulfment of AIEC-LF82 within the gut epithelium

Interactions of the invading microbe with the host cellular processes is a key trigger for the generation of inflammatory responses. Although phagocytic cells are primarily engaged in the uptake and clearance of microbes, it is well known that microbes do enter through epithelial TJs^37^. What is unknown is whether the epithelial cell relies on the engulfment pathway for uptake and subsequently clear them via the phagolysosomal pathway. To understand the role of ELMO1 during bacterial entry into epithelial cells, we used EDMs generated from colons of Wild type (WT) and ELMO1^−/−^mice. Depletion of ELMO1 in the EDMs from ELMO1^−/−^mice was confirmed by immunoblotting (Figure 4*A*). When bacterial internalization was measured using the well-accepted gentamicin protection assay^18, 38^, we found that compared to WT controls (n = 10), internalization at 1 h after infection was decreased by 73% decrease in ELMO1 KO EDMs (n = 8) (*p* < 0.05; Figure 4*B*).

**Figure 4.**
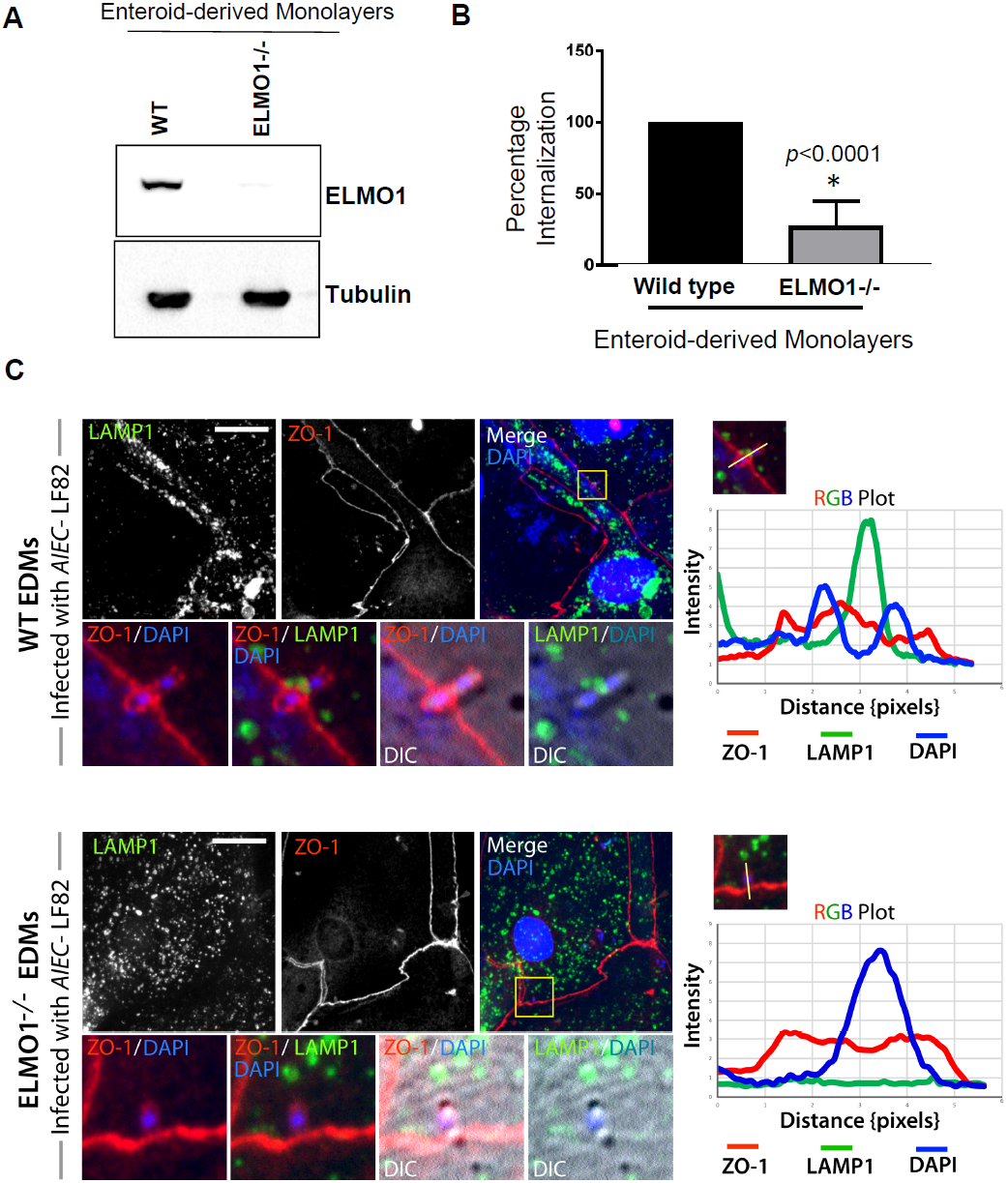
The engulfment (internalization) of *AIEC*-LF82 through epithelial TJs is impaired in ELMO1^−/−^EDMs with reduced recruitment of lysosomal proteins to the sites of internalization. *(A)* Expression of ELMO1 protein was assessed by immunoblotting in enteroids isolated from colons of WT and ELMO1**-/-**mice. a-Tubulin was analyzed as a loading control. *(B)* WT and ELMO1^−/−^EDMs were infected with *AIEC*-LF82 for 3 h prior to assessment of bacterial internalization using gentamicin protection assay (see *Methods*). Bar graphs display % internalization. Data represent the mean ± SD of three separate experiments. * indicates p≤0.05 as assayed by two-tailed Student’s t test. *(C)* WT and ELMO1^−/−^EDMs were infected with *AIEC*-LF82 as in *B*, fixed, stained with ZO1 (red), LAMP1 (green) and DAPI for nucleus, and analyzed by confocal imaging. *<u>Left</u>*: Maximum projection of Z-stacks of representative fields were shown. Insets in merged images represent magnified images and displayed at the bottom to zoom in at the point of bacterial entry through epithelial TJs. Lysosomes (marked by LAMP1) were aligned with the TJs (marked by ZO-1) in WT EDMs, but remain dispersed throughout the epithelial cell in ELMO1^−/−^EDMs. Lysosomes were seen in close proximity to the invading bacteria exclusively in the WT EDMs. *<u>Right</u>*: RGB plots show distance in pixels between the internalized bacteria (blue) and the TJs of host cells (red) and lysosomes (green).

**Figure 5:**
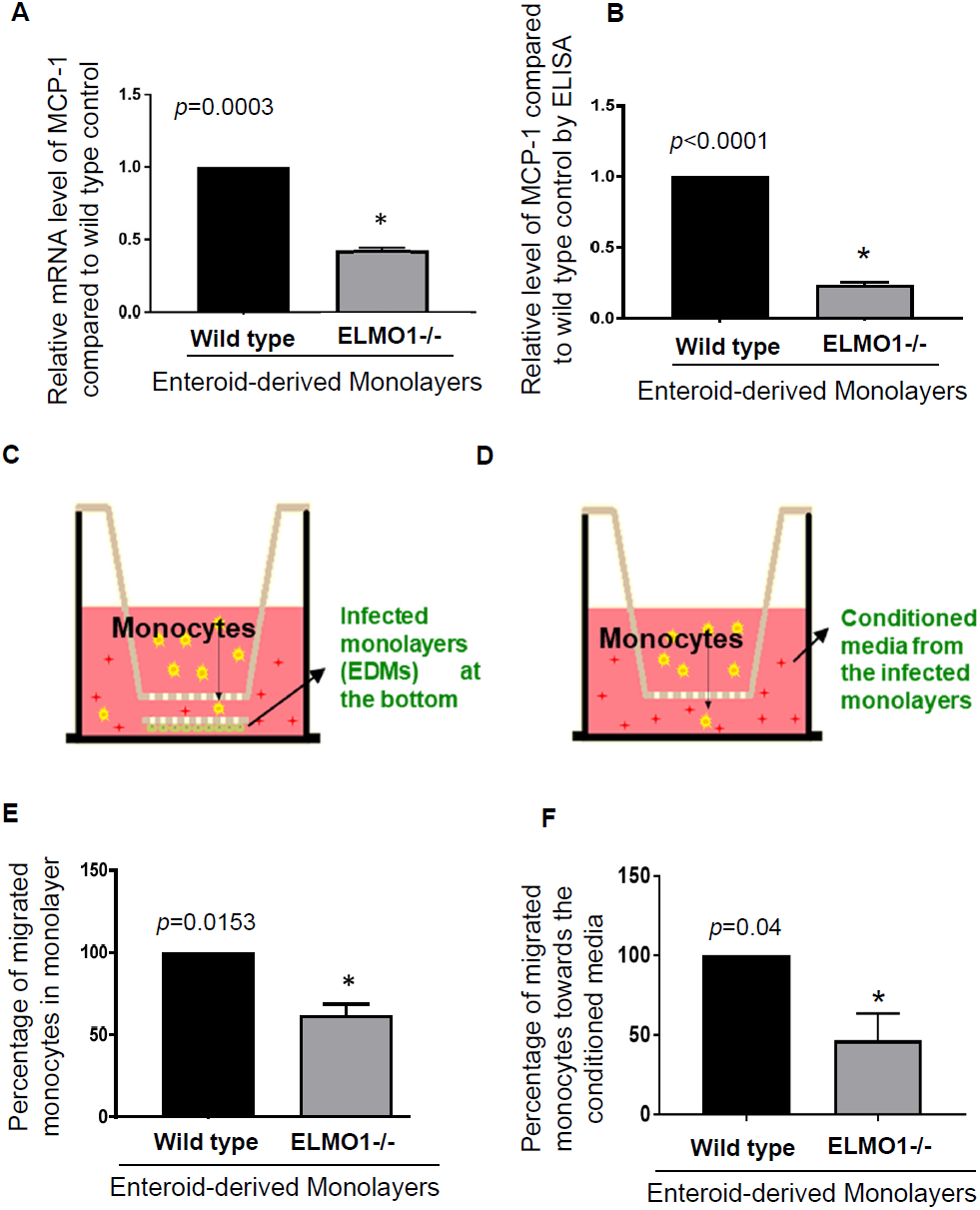
The induction of MCP-1 and recruitment of monocytes in response to *AIEC*-LF82 is blunted in ELMO1^−/−^EDMs; compared to WT EDMs. *(A)* Levels of expression of MCP-1 was measured by qRT-PCR in EDMs derived from WT and ELMO1^−/−^mice after infection with *AIEC*-LF82 for 6 h. Bar graphs display fold difference in MCP-1; mean ± SD of three separate experiments. * indicates p≤0.05 as assayed by two-tailed Student’s t test. *(B)* Infection-induced production of MCP-1 by WT and ELMO1^−/−^ EDMs in **A** was also measured by ELISA on the supernatant collected after 6 h post-infection. Data represent the mean ± SD of three separate experiments. *(C-D)* Schematics of the EDM-monocyte coculture model used to study monocyte recruitment. Either infected EDMs (WT or ELMO1^−/−^) *(C)* or conditioned supernatant *(D)* collected from infected EDMs was placed in the lower compartment separated from monocytes (upper chamber) separated by porous inserts of Transwell™ (see *Methods*). The number of monocytes that migrated from the upper to the lower chamber by 12 h was counted. *(E-F)* Bar graphs display monocyte migration towards infected EDMs *(E)* or conditioned media *(F)* plotted as percent (%) normalized to that seen when using supernatant from WT EDMs. Data represent as mean ± SD of three separate experiments. * indicates p≤0.05 as assayed by two-tailed Student’s t test.

We also carried out confocal immunofluorescence studies to evaluate how the bacteria enter the epithelial cells. We found that in both WT and ELMO1^−/−^EDMs the *AIEC*-LF82 enter through the epithelial TJs, as determined by ZO-1 staining that surrounded the bacteria (Figure 4*C*). However, we found dissimilarities when it comes to the proximity of lysosomes to the invading pathogens. In WT EDMs, lysosomes (as detected using the lysosomal integral membrane protein, LAMP1) were found in close proximity to the invading *AIECs* (Figure 4*C*) in WT EDMs, indicating that lysosomes are recruited to the site of TJ breach; such approximation was not seen in the ELMO1^−/−^EDMs. These findings raise the possibility that in the absence of lysosome targeting, ELMO1^−/−^EDMs may be defective not just in bacterial uptake, but also in bacterial clearance. These findings are consistent with our previously published role of ELMO1 in the clearance of another invasive pathogen, *Salmonella*^17^. Although typically longer time points are required to assess clearance, such assays were not possible using EDMs as model system. This is because one of the major limitations of EDMs is that when infected, they typically undergo extensive cell death within 8 - 12 h, making any cellular morphological analysis technically impossible.

Taken together, our results using ELMO1^−/−^EDMs demonstrate that ELMO1 is required for *AIEC*-LF82 uptake through breaches in the epithelial TJs, and for the proper targeting of lysosomes to the invading *AIEC*-LF82 pathogen. Morphologic findings in EDMs predict that once internalized, ELMO1 may also be required for efficient clearance of the *AIEC*-LF82 (studied in detail with macrophages in Figure 6*B* and Supplementary Figure 1).

**Figure 6:**
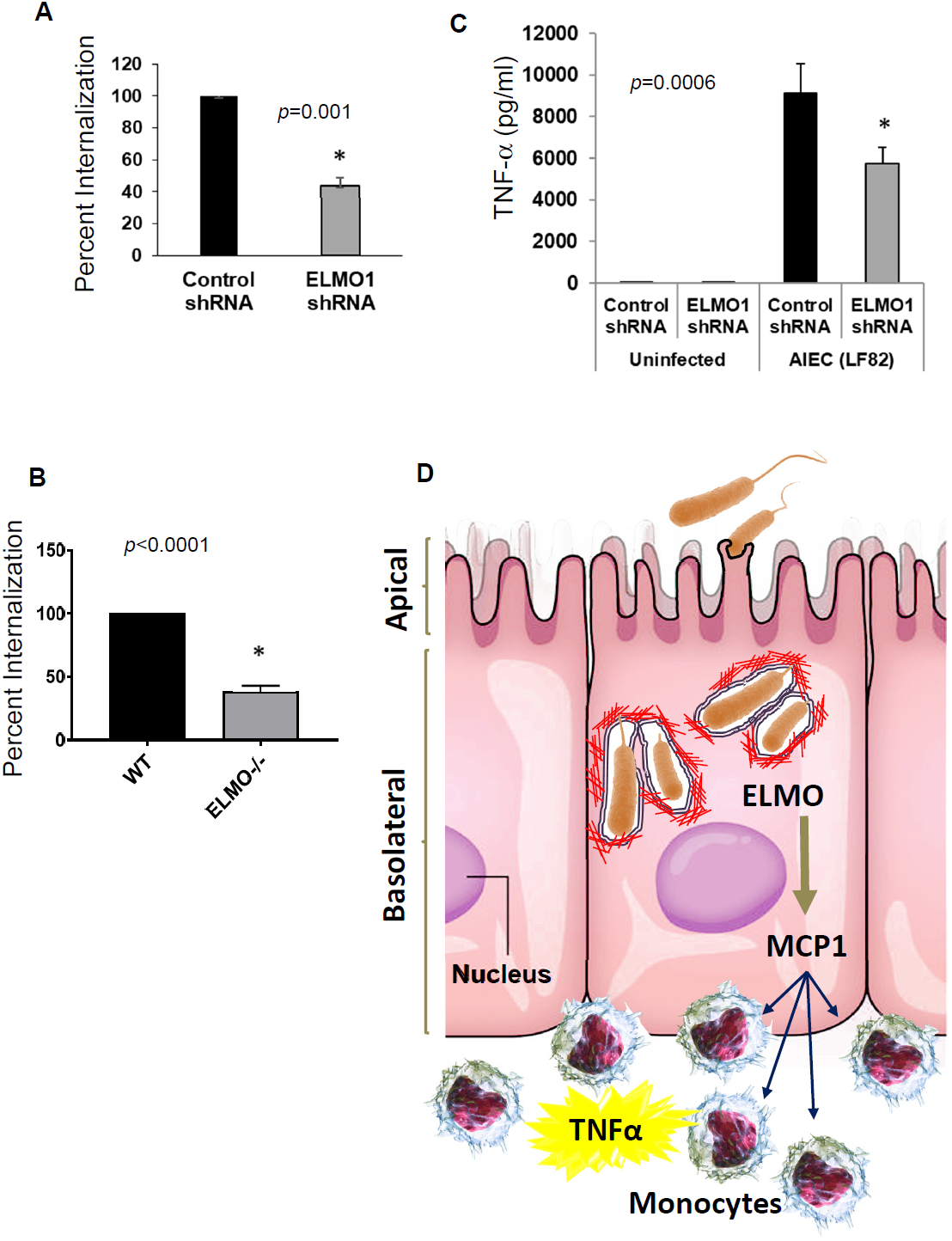
Compared to WT macrophages, ELMO1-deficient macrophages display an impairment in the engulfment of *AIEC*-LF82 and induction of TNF-α. *(A)* Internalization of *AIEC*-LF82 in control (Control shRNA) and ELMO1-depleted (ELMO1 shRNA) J774 cells was assessed using gentamicin protection assay as in 3*B*. Bar graphs display % internalization observed at 3 h after infection. Findings are represented as mean ± SD of three separate experiments, normalized to Control shRNA. * indicates p≤0.05 as assayed by two-tailed Student’s t test. *(B)* The intestinal macrophages isolated from wild type (WT) and ELMO1 -/-mice were infected with *AIEC*-LF82 for 1 h at 37°C and internalization was measured by the gentamicin protection assay. The average number of internalized bacteria (mean ± SD) was calculated and represented as % internalization. *(C)* TNF-a produced by *AIEC*-LF82**-**infected J774 cells in A were analyzed by ELISA with the ELISA after 3 h of infection. Data represent as mean ± SD of three separate experiments. * indicates p≤0.05 as assayed by two-tailed Student’s t test. *(D)* Schematic summarizing the role of ELMO1 in coordinating inflammation first in non-phagocytic (epithelial) and subsequently in phagocytic (monocytes) cells of the gut. Epithelial ELMO1 is essential for the engulfment of invasive pathogens like *AIEC*-LF82 and for the induction of MCP-1 in response to such invasion. MCP-1 produced by the epithelium triggers the recruitment of monocytes, facilitating their recruitment to the site of infection. Once recruited, ELMO1 in monocytes is essential for the engulfment and clearance of invasive bacteria and for the production of pro-inflammatory cytokines such as TNF-α. MCP-1 and TNF-α released from the epithelial and monocytic cells initiates a chain reaction for the recruitment and subsequent activation of other monocytes and T-cells. The resultant storm of pro-inflammatory cytokines propagates diseases characterized by chronic inflammation.

### 3.4. ELMO1 in the gut epithelium is required for the generation of pro-inflammatory cytokine MCP-1 following infection with AIEC LF82

We previously showed that ELMO1 and an intact host engulfment pathway is essential for the induction of pro-inflammatory cytokine MCP-1 in monocytes^15^. Because the inflamed gut epithelium can express MCP-1^39^, and because MCP-1 plays a major role in recruiting monocytes that in turn generates inflammatory cytokines in CD-afflicted gut^39-42^, we hypothesized that ELMO1 may be required also for MCP-1 production by the gut epithelium once it is breached by invading *AIEC*-LF82.

We first asked if there is any correlation between ELMO1 expression and MCP-1 in CD-derived EDMs. To this end, we measured the levels of MCP-1 by qRT-PCR (Figure 5*A*) and by ELISA (Figure 5*B*). In WT EDMs, MCP-1 was undetectable without infection, but its levels were elevated after infection. Compared to WT EDMs, infection-triggered induction of MCP-1 was blunted in ELMO1^−/−^ EDMs (Figure 5*A-B*).

Taken together, these findings demonstrate that ELMO1 is required for the generation of MCP-1 by the epithelium that is breached by CD-associated invasive pathogens, and that the ELMO1-MCP-1 axis may be a fundamental pathway that responds to dysbiosis in the gut lumen.

### 3.5. An intact ELMO1-MCP-1 axis in the gut epithelium is required for the recruitment of monocytes to the infected epithelium

Previous studies have demonstrated that CCL2/MCP-1^−/−^mice had significant reduction in monocyte recruitment in inflammatory models and Th2 cytokines (IL-4, IL-5 and IFN-γ) in the secondary pulmonary granulomata in response to *Schistosoma mansoni* eggs^43, 44^. To understand the role of the ELMO1-MCP-1 axis in the recruitment of monocytes, we infected the WT and ELMO1^−/−^EDMs with *AIEC*-LF82 for 6 h and assessed the ability of these EDMs to recruit monocytes. After the infection, either the conditioned supernatant (Figure 5*C*) or the infected monolayer itself (Figure 5*D*) was co-cultured with monocytes, and migration of monocytes toward the infected EDM site was assessed. Compared to WT EDMs, ELMO1^−/−^EDMs displayed a 50% reduction in monocyte recruitment. These results indicate that ELMO1-dependent MCP-1 production by the gut epithelium could serve as an upstream cue for monocyte recruitment to the sites of infection.

### 3.6. ELMO1 in macrophages is essential for the engulfment of AIEC-LF82, and for the generation of pro-inflammatory cytokines

Next, we determined how ELMO1 impacts macrophage response upon being recruited to the sites of *AIEC*-LF82 infection. To study the role of ELMO1 in the internalization of *AIEC*-LF82, we used the gentamicin protection assay to assess bacterial uptake in ELMO1-depleted J774 macrophages^15^ (ELMO1 shRNA; around 90% depletion confirmed by immunoblotting). ELMO1-depleted cells showed approximately 50% reduction in bacterial internalization compared to WT cells (p value 0.001; Figure 6*A*).

To determine whether ELMO1 is essential for the clearance of *AIEC*-LF82 (as observed in *Salmonella*^17^), bacterial engulfment and clearance was studied at 30 min, 3, 6, 12 and 24 h in the murine macrophage cell line J774 (Supplementary Figure 1). ELMO1^−/−^cells showed lower uptake of *AIEC*-LF82 compared to WT macrophages (50% reduction; *p value 0.001*) at 30 min, but retention of a higher bacterial load at later time points (3 fold increase at 24 h after infection; with a *p value 0.05*). These findings indicate that ELMO1 is required not just for uptake, but also for clearance of *AIEC*-LF82.

To assess the contribution of ELMO1 in bacterial internalization in a more physiologically relevant system, we carried out the same assay as above, but replaced of J774 cultured cell lines with primary intestinal macrophages enriched from WT or ELMO1^−/−^mice (Figure 6*B*). While intestinal macrophages from WT mice engulfed bacteria efficiently, bacterial uptake was decreased approximately 60% in macrophages from ELMO1^−/−^mice (*p value <0.0001*), indicating that ELMO1 is essential for the engulfment of *AIEC*-LF82 in macrophages.

TNF-α is a major pro-inflammatory cytokine that is elevated early in the development of CD^39, 45^. Macrophages exposed to *AIECs* engulf the bacteria, and can induce TNF-α^39^. We analyzed how reduced engulfment in the absence of ELMO1 impacts the release of TNF-α into the supernatant from control and ELMO1-depleted [by shRNA] J774 macrophages that were infected with *AIEC*-LF82. Using ELISA to detect the cytokine, we found that, the ELMO1-depleted macrophages had significant reduction in TNF-α compared to control shRNA cell (*P value 0.0006*; Figure 6*C*). These findings are consistent with our previous findings in the context of *Salmonella*^15^.

## Discussion

The major finding of our work is the identification of the engulfment pathway that is coordinated by ELMO1 as a novel host response element, which operates in two different cell types in the gut, i.e., the epithelium and the monocytes. ELMO1 facilitates the engulfment of pathogenic microbes in the gut epithelium, triggers the induction of the pro-inflammatory cytokine, MCP-1; the latter helps recruit monocytes from peripheral blood to the site of local inflammation (Figure 6*D*). Our results also highlight and exemplify the complex multi-step interplay between luminal dysbiosis, which is an invariant hallmark in IBD and macrophages, key players in the innate immune system of the gut. First, we show that the pathogenic *AIEC*-LF82 strain that is associated with CD can invade the gut epithelial lining by entering through epithelial TJs, and subsequently triggers the production of the pro-inflammatory cytokine, MCP-1 before being cleared via the phagolysosomal pathway. In the absence of ELMO1, uptake of the bacteria into the epithelial cells is impaired, and MCP-1 production is blunted. This ELMO1-MCP-1 axis then triggers the recruitment of monocytes. Next, we showed that the same molecule, ELMO1 is also essential for the uptake and clearance of *AIEC*-LF82 in the monocytes, and is required for coordinately mounting yet another pro-inflammatory cytokine response, TNF-α. This ELMO1-TNF-α axis presumably feeds forward to propagate inflammation in the gut by triggering the activation of other monocytes and T cells. Thus, the two signaling axes, ELMO1-MCP-1 and ELMO1-TNF-α, orchestrated by the same engulfment pathway in two different cell types, the epithelium and the monocytes, respectively, appear to be working as ‘first’ and ‘second’ responders to combat pathogenic microbes, thereby relaying distress signals from one cell type to another as the microbe invades through the breached mucosal barrier.

Until now, the majority of IBD-related research and therapeutic strategies have remained focused on T cell responses and on neutralizing the impact of TNF-α. By demonstrating the presence of two hierarchical spatially and temporally separated signaling axes, our work provides mechanistic insights into some of the upstream/initial immune responses that play out in the epithelium and within the macrophages upon sensing luminal dysbiosis. This 3-way interaction between microbe-epithelium-macrophages is crucial to maintain homeostasis, and intestinal macrophages maintain the balance between homeostasis and inflammation^46^. A breach in the epithelium brought about by invading pathogens shifts the balance towards pro-inflammatory pathways. This work defines an upstream event that could be exploited to develop biomarkers, and eventually interrogated for the identification of strategies for therapeutic intervention (e.g., anti-MCP-1 therapy, as discussed later). The need for an in-depth understanding of the nature and the extent of the contribution of epithelial cells and/or monocytes in disease progression is urgent because of the limited efficacy of the available treatment options^47^. Because the recruitment of monocytes from circulation to the site of infection/inflammation is a key early event in inflammatory diseases of the gut, the ELMO1→MCP-1 axis we define here is potentially an actionable high value diagnostic and therapeutic target in IBD. Detection of high levels of ELMO1 in the epithelium could serve as an early indicator of activation of the engulfment pathway, and hence, could serve as a surrogate diagnostic marker of early inflammation due to luminal dysbiosis. Similarly, targeting the engulfment pathway is expected to restore immune homeostasis and resolve chronic inflammation via a completely novel approach that could synergize with existing therapies, and thereby, improve response rates and rates of sustained remission.

This work also provides the first mechanistic insights into how luminal dysbiosis initiates inflammation in the gut. Bacterial clearance and microbial dysbiosis are one of the hallmark of CD that control the outcome of innate immune responses. Healthy commensals like *Bacteroidetes* and *Faecalibacterium prausnitzii* are decreased in patients with CD, while pathogenic microbes like invasive *Escherichia coli, Serratia marcescens, Cronobacter sakazakii* and *Ruminoccus gnavus* are increased ^9,10,14,48^. A dysbiotic microbial population can harbor pathogen and pathobionts that can aggravate intestinal inflammation or manifest systemic disease. Effector proteins produced by pathogenic bacteria can activate signaling that induce granuloma formation; one of the key symbol in CD pathogenesis^49^. In CD granuloma, the number of mucosal adherent invasive *E. coli* is higher because of defective clearance and that can cause dysbiosis^8, 50^. Previously we showed that ELMO1 and the engulfment-pathway in the professional phagocytes is essential for the internalization of *Salmonella,* and for mounting intestinal inflammation^15^. In this work, we have demonstrated the role of ELMO1 in non-phagocytic cells in the context of IBD using the CD-associated *AIEC-LF82*. The use of stem-cell based enteroids from ELMO1-/-mice, either alone or in cocultures with monocytes allowed us to interrogate the function of ELMO1 in the epithelium and the monocytes separately. However, the complete impact of ELMO1 in chronic inflammatory diseases need further investigation.

Finally, by showing that the ELMO1-MCP-1 axis is an early step in gut inflammation, this work provides rationale and impetus for the development of anti-MCP1 biologics to treat IBD. MCP-1 belongs to a CC chemokine subfamily, and its effects are mediated through CC chemokine receptor 2 (CCR2). So far, in human, only A2518G variation in MCP-1 gene promoter has been associated with CD^51^. However, MCP-1 is not just important in IBD, but also involved in other inflammatory diseases, such as atherosclerosis^52^. In fact, MCP-1 promotes the balance between anti-inflammatory and pro-inflammatory responses to infection. Treatment with recombinant MCP-1/CCL2 increases bacterial clearance and protects mice that are systemically infected with *Pseudomonas aeruginosa* or *Salmonella typhimurium*^53^. Administration of MCP-1 can increase chemotaxis on murine macrophages, enhance phagocytosis and killing of bacteria^53^, whereas pretreatment of mice with anti-MCP-1/CCL2 impaired bacterial clearance. Therefore, increased expression of MCP-1 by ELMO1 in intestinal epithelium after exposure to *AIEC*-LF82 is likely to have a two-fold importance--1) for controlling the increased bacterial load by killing the bacteria, and 2) for promoting monocyte recruitment and activation, which initiate a pro-inflammatory cytokine storm by inducing TNF-ɑ from macrophages. Further work is required before we can begin to assess the safety and efficacy of anti-MCP-1 therapies in IBD.

## Acknowledgement

This work was supported by NIH grants DK107585, DK099275; NIH CTSA grant UL1TR001442 (to S.D), CA100768, CA160911 and DK099226 (to P.G). S.R.I was supported by NIH Diversity Supplement award (3R01DK107585-02S1) and Y.M was supported by NIH Training Grant in Gastroenterology (T32DK0070202). We are thankful to Mitchel Lau and Julian Tam for technical support. We are grateful to Drs. Ernst, Crowe, Barrett and Eckmann for providing the access to the shared equipment within the UCSD GI division.

## Isolation of enteroids from colonic specimens of mouse and human

The crypts after digestion with Collagenase were filtered with a cell strainer and washed with medium (DMEM/F12 with HEPES, 10% FBS), After adding collagenase I solution containing gentamicin (50 mg/ml, Life Technologies) and mixing thoroughly, the plate was incubated at 37° C inside a CO2 incubator for 10 min, with vigorous pipetting between incubations and monitoring the intestinal crypts dislodging from tissue. The collagenase was inactivated with media and filtered using a 70-μm cell strainer over a 50-ml centrifuge tube. Filtered tissue was spun down at 200 g for 5 min and the media was aspirated. The epithelial units were suspended in matrigel (BD basement membrane matrix). Cell-matrigel suspension (15 ml) was placed at the center of the 24-well plate on ice and placed on the incubator upside-down for polymerization. After 10 min, 500 μl of 50% conditioned media (prepared from L-WRN cells with Wnt3a, R-spondin and Noggin, from the laboratory of Thaddeus S. Stappenbeck^1^) containing 10 μM Y27632 (ROCK inhibitor) and 10 μM SB431542 (an inhibitor for TGF-β type I receptor) were added to the suspension. For the human colonic specimens Nicotinamide (10 μM, Sigma-Aldrich), N-acetyl cysteine (1 mM, Sigma-Aldrich), and SB202190 (10 μM, Sigma-Aldrich) were added to the above media. The medium was changed every 2 days and the enteroids were expanded and frozen in liquid nitrogen.

### RNA Preparation, Real-Time Reverse-Transcription Polymerase Chain Reaction

RNA was prepared using RLT buffer (Qiagen Beverley, Inc) and β-mercaptoethanol. Total RNA was extracted using the RNeasy Microkit (Qiagen Beverly, Inc) and reverse transcribed with a cDNA Supermix (Qiagen Beverly, Inc), both according to the manufacturer’s instructions and as done previously^2, 3^. Real-time RT PCR was performed using SYBR Green High ROX (Biotool) with primers (Intergrated DNA Technologies) detected using StepOnePlus Real-Time PCR Systems (Applied Biosystems) and normalized to the values of β-actin for mice and GAPDH for human. The fold change in mRNA expression was determined using the ΔΔCt method as done previously^2-4^.

### PCR primers

Mouse CCL2 (MCP-1), KC (IL-8), and β-actin mRNA were amplified by using the primers: CCL2-F 5’ aagtgcagagagccagacg 3’ and CCL2-R 5’ tcagtgagagttggctggtg 3’; KC-F 5’ cctgctctgtcaccgatgt 3’ and KC-R 5’ cagggcaaagaacaggtcag 3’.

Human CCL2 (MCP-1), ELMO1, IL-1β, TNF-α, and GAPDH mRNA were amplified by using the primers: CCL2-F 5’ agtctctgccgcccttct 3’ and CCL2-R 5’ gtgactggggcattgattg 3’; ELMO1-F 5’ cagaagatcatgaagccttgc 3’ and ELMO1-R 5’ tccatcagctcaacgaagg 3’; TNF-α-F 5’ cgctccccaagaagacag 3’ and TNF-α-R 5’ agaggctgaggaacaagcac 3’.

### Gene expression analysis

This cohort included gene expression data from multiple publicly available NCBI-GEO data-series (10714, 10961, 11831, 12945, 13067, 13294, 13471, 14333, 15960, 17538, 18088, 18105, 20916, 2109, 2361, 26682, 26906, 29623, 31595, 37892, 4045, 4107, 41258, 4183, 5851, 8671, 9254, 9348), and contained information on 214 unique normal colon samples. All 214 samples contained in this subset were cross-checked to exclude the presence of redundancies/duplicates. To investigate the relationship between the mRNA expression levels of selected genes (i.e. ELMO1 and CCL2), we applied the *Hegemon*, “*hierarchical* exploration of *gene expression* microarrays on-line” tool ^5^. The Hegemon software is an upgrade of the BooleanNet software^6^, where individual gene-expression arrays, after having been plotted on a two-axis chart based on the expression levels of any two given genes, can be stratified using the StepMiner algorithm and automatically compared for statistically significant differences in expression. We stratified the patient population of the NCBI-GEO discovery dataset in different gene-expression subgroups, based on the mRNA expression levels of ELMO1. Once grouped based on their gene-expression levels, patient subsets were compared for CCL2.

### Confocal Microscopy

After 1 h infection, media was aspirated, and cells were treated with 5% CM media with 250 μg/ml gentamicin for 90 min. Media was aspirated and 5% CM media was added to the wells. After 6 h of total infection time, samples were washed in 1-X PBS, pH 7.4 and fixed in 2% formaldehyde, washed with PBS, and permeabilized with 0.1% saponin-2 % BSA (Sigma-Aldrich) in PBS for 10 minutes. Cells were blocked with 0.05% saponin-1 % BSA in PBS (blocking solution) subsequently incubated with LAMP1 (Biolegend) and ZO1 (Santa Cruz cat#sc-33725) overnight in blocking solution, diluted 1:800 and 1:140 respectively. The secondary antibodies, goat anti-mouse-Alexa488 (Life technologies, cat#A-11017 1/500), goat anti-rabbit-Alexa594 (Life technologies, cat#A-11012 1/500) and DAPI (1/1,000) were prepared in blocking solution. Images were acquired using a Leica TCS SPE CTR4000 / DMI4000B-CS Confocal microscope with a Plan APO 63x objective. Multi-color images were obtained using excitation laser lines 405, 488 and 543 and transmission light, with respective detection. Z-stack acquisition was performed using a 1024×1024 pixels (58.3×58.3 micron) with a total of 10 sections (0.35 micron thickness). Images were analyzed in FIJI of ImageJ. Image flattening was obtained using average projection.

### Isolation of murine intestinal macrophages

Murine intestinal macrophages were isolated using methods described previously^2,3,7^. Briefly, small intestines were collected and intestinal fragments were digested with enzyme cocktail including collagenase (Miltenyi Biotec). The cell suspension was passed through a cell strainer to remove debris, and enriched with CD11b^+^cells using the CD11b MACS kit (Miltenyi Biotec). Intestinal macrophages were labeled using monoclonal antibody CD11b (BD Biosciences) and sorted using a FACS Vantage (BD Biosciences). The purity of intestinal macrophages were determined by staining with antibody specific for the macrophage marker F4/80.

---

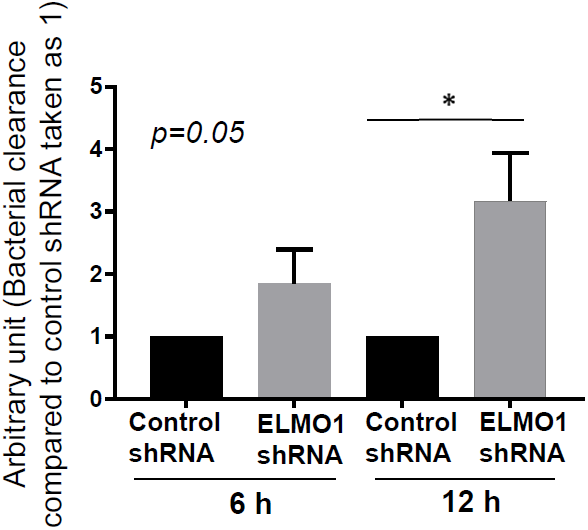

